# A sinusoidal transform of the visual field in cortical area V2

**DOI:** 10.1101/2020.12.08.416651

**Authors:** Madineh Sedigh-Sarvestani, Kuo-Sheng Lee, Rachel Satterfield, Nicole Shultz, David Fitzpatrick

## Abstract

The retinotopic maps of many visual cortical areas are thought to follow the fundamental principles that have been described for primary visual cortex (V1) where nearby points on the retina map to nearby points on the surface of V1, and orthogonal axes of the retinal surface are represented along orthogonal axes of the cortical surface. Here we demonstrate a striking departure from this conventional mapping in the secondary visual area (V2) of the tree shrew. Although local retinotopy is preserved, orthogonal axes of the retina are represented along the same axis of the cortical surface, an unexpected geometry explained by an orderly sinusoidal transform of the retinal surface. This sinusoidal topography is ideally suited for achieving uniform coverage in an elongated area like V2, is predicted by mathematical models designed to achieve wiring minimization, and provides a novel explanation for stripe-like patterns of intra-cortical connections and stimulus response properties in V2. Our findings suggest that cortical circuits flexibly implement solutions to sensory surface representation, with dramatic consequences for the large-scale layout of topographic maps.

## Introduction

The discovery of retinotopic maps, the orderly representation of the visual field in cortical and subcortical structures, has had enormous impact on our understanding of the visual system and its development (*1, 2*). Classic descriptions of retinotopic maps emphasize the spatial preservation of local and global retinal relationships as a linear conformal transform in which: (1) orthogonal axes of the visual field are mapped along orthogonal axes of the cortical surface and (2) progression across the map yields smooth and continuous unidirectional progression in the visual field with no reversals. Moreover, this linear transform organization is predicted by computational models that achieve optimal coverage of visual space while minimizing wiring costs (*3*–*5*). Primary visual cortex (V1) in a number of species has been shown to adhere to these principles (*6*–*10*) and although deviations have been noted in higher visual areas (*5, 11, 12*), no alternate framework has been proposed. Here we present evidence from studies of the secondary visual area (V2) of the tree shrew--a close relative of primates with a smooth brain that is ideal for *in vivo* imaging--for a dramatic departure from the classic conformal framework of retinotopic maps.

Using calcium imaging in awake animals, we found that the transformation of the visual field in area V2 is sinusoidal, with representation of the two orthogonal dimensions of the visual field along the same elongated dimension of V2. Mathematical models that simulate the formation of cortical maps show that the same principles that give rise to linear conformal maps in V1 give rise to sinusoidal maps in V2, and that the difference in mapping can be explained by different cortical area geometries. Moreover, we show with anatomical tracing studies that the organization of feed-forward connections from V1 is sufficient to build the sinusoidal map in V2 from the linear map in V1. Aside from its consequence on retinotopic maps, this intricate wiring also influences the organization of other functional feature maps, producing a stripe-like pattern reminiscent of primate V2 (*13*–*15*).

Altogether, our study reveals a novel transform of the retinal surface on the cortex, which arises from precise V1 connectivity to achieve optimal visual field coverage in V2, and is correlated with the large-scale organization of functional feature maps. This suggests that hierarchical connectivity between visual areas can give rise not only to increasing complexity in receptive field properties, but also to increasing complexity in large-scale retinotopic organization. More importantly, our findings indicate that linear mapping of sensory receptor surface is just one solution to optimal surface coverage and that cortical wiring can flexibly produce drastically different solutions under different conditions.

## Results

To characterize the retinotopic maps in L2/3 of V1 and V2, we used 1-photon widefield and 2-photon single-cell fluorescence microscopy to image neuronal calcium dynamics (AAV9.Syn.GCaMP6s.WPRE.SV40) in awake head-fixed and restrained tree shrews trained to remain motionless during recordings. Retinotopic maps in adjacent regions of V1 and V2 were imaged simultaneously using a 3-5 mm cranial window implanted over the border of V1 and V2, covering the full ∼1.5 mm width of V2 along with a smaller fraction of V1 medial to the V1/V2 border (Figure 1A). The medial and lateral border of V2 were defined based on the retinotopic map (Methods). To quantify the retinotopic transform, we used stimuli (see Methods) that varied systematically in position along the elevation (vertical) and azimuth (horizontal) axes of visual space.

**Figure 1:**
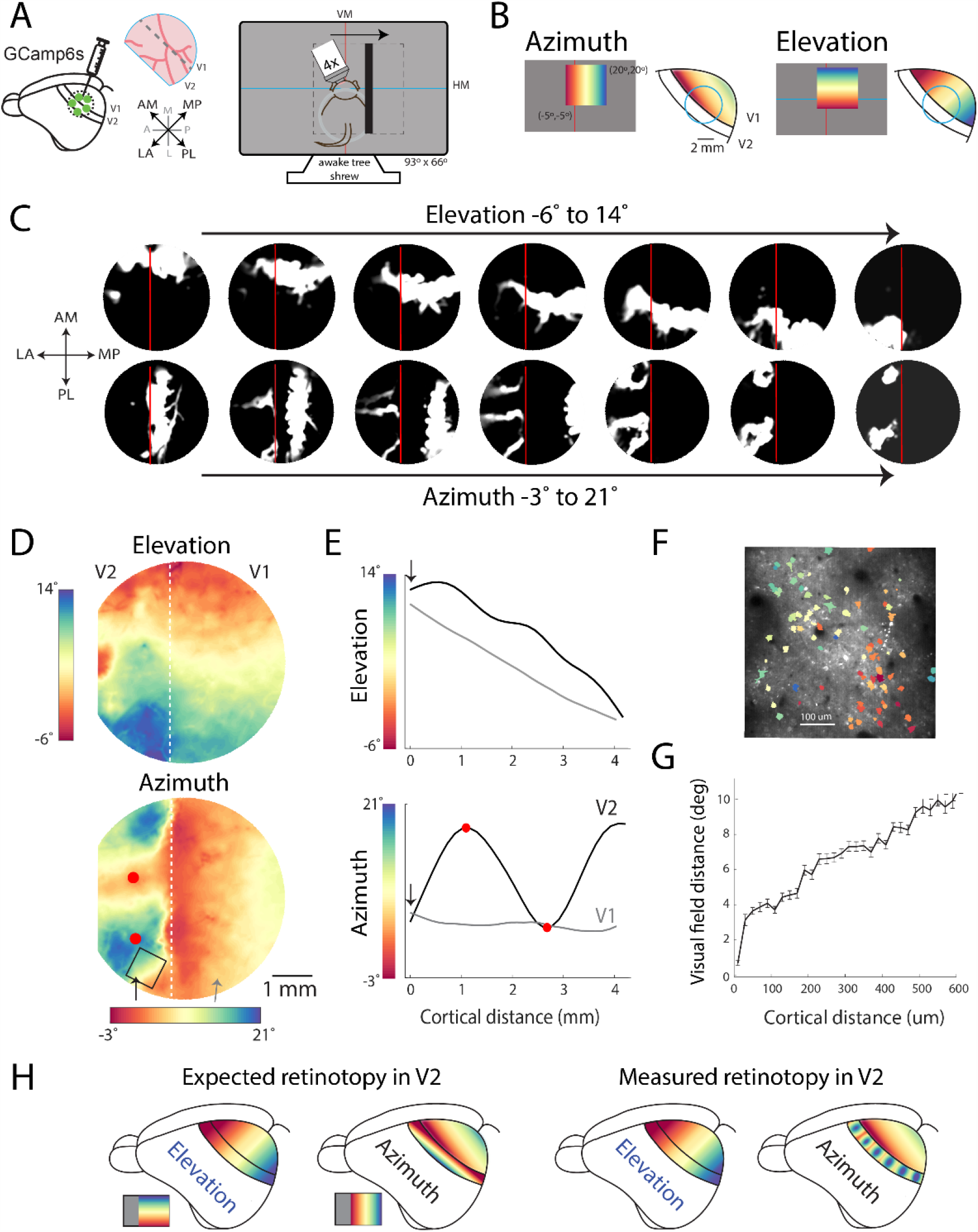
Tree shrew V2 exhibits a sinusoidal transform of the visual field. **A)** Calcium indicator was injected over V1 and V2. **B)** Animals viewed stimuli varying in azimuth or elevation. Schematized retinotopic maps of V1. **C)** Magnitude of trial-averaged calcium response during equal intervals of bar position, thresholded at mean ± 0.3 SD to highlight spatial extent. **D)** Azimuth and elevation maps (Methods). **E)** Extracted azimuth, and elevation values corresponding to two straight lines in V1 and V2. Red dots indicate gradient reversals. **F)** Two-photon FOV and single-cell ROIs colored by preferred azimuth. **G)** Pairwise distance in visual field along azimuth vs. cortical distance for cells in F. Error bars are SEM. **H)** Unlike the expected orthogonal maps in V2, azimuth is also mapped along the long dimension with nearby regions alternately representing more central or peripheral regions of the visual field.

Consistent with previous studies (*10, 16*) of retinotopy in tree shrew V1 (Figure 1B) we found that the elevation dimension of the contralateral visual field was mapped smoothly along the AM-PL axis of V1 and the azimuth dimension was mapped smoothly along the orthogonal LA-MP axis (Figure 1 C, D. Note that for simplicity in this and remaining figures, we have rotated the cranial window clockwise ∼45 degrees so that elevation and azimuth dimensions in V1 are roughly vertical and horizontal respectively.

The map of elevation in area V2 appeared similar to the map in the adjacent portion of V1, a relatively smooth progression with lower portions of the visual field represented by more anterior portions of cortex (Figure 1 C, D). In contrast, the representation of the azimuth dimension in V2 was unlike any retinotopic map that has been described. We had expected that the azimuth map would be organized orthogonal to the elevation map such that a vertical (iso-azimuth) bar in visual space would give rise to a continuous band of activity in V2 parallel to the V1/V2 border and that this pattern would progress away from the V1/V2 border with eccentricity, as a mirror image to what was present in V1 (Figure 1H, left).

Instead, a single vertical bar activated multiple distinct narrow bands that extended across the width of V2 (Figure 1C). The distance between the bands changed systematically as a function of the bar’s position in visual space, merging for stimuli at the most central and peripheral represented locations.

The striking nature of this unusual mapping becomes evident in the pseudo-color map of azimuth (Figure 1D) which shows azimuth represented in an oscillatory fashion along the AM-PL axis of V2, with alternating stripes sensitive to relatively more central or peripheral portions of the visual field. The oscillatory nature of this pattern is best appreciated by plotting the azimuth values corresponding to a linear trajectory in V2 (Figure 1D arrowheads). Such a path on the cortex reveals a linear mapping of azimuth in V1 but a sinusoidal mapping in V2 (Figure 1E). The peaks and valleys of the azimuth oscillations lie at the center of the striped regions (red dots) in the azimuth map of V2 and correspond to ‘reversals’ in the orderly progression in azimuth or the ‘merger’ of two opposing gradients in mapping. The period of these oscillations, from the most central to most peripheral point, was 1.25+/-0.35 mm (n=6 animals, Methods). Oscillating patterns for azimuth were found in every animal, albeit with variations in shape (Figure S1).

Our epifluorescence imaging results suggest that this large-scale oscillatory azimuth pattern occurs with smooth changes in the visual field trajectory, but we thought it was important to confirm this with 2-photon cellular level resolution. We focused on regions in V2 with the highest rate of change in azimuth, the region between central and relatively more peripheral stripes (Figure 1D box). Consistent with our wide-field data, single-cell measurements of visual field sensitivity along the azimuth axis (Figure 1F) showed smooth and continuous changes in azimuth location as a function of cortical distance between cell pairs (Figure 1G).

These results imply a fundamentally different transform for the mapping of retinotopy in V2 from what has been described in V1 and in most other visual areas (Figure 1H). A better understanding of this transform requires considering the fine-scale relationships between the maps of azimuth and elevation. We defined the spatial relationships between iso-elevation and iso-azimuth contours corresponding to equal distances in the visual field (Figure 2A). Consistent with previous studies, in V1, iso-azimuth and iso-elevation contours in the tree shrews (*10*) and other animals (*6*–*9*) exhibit a strong bias for orthogonality (Figure 2 B, C). Just as elevation and azimuth contours are orthogonal in the visual field, they are orthogonal in V1 (Figure S2A) : i.e., the transformation of the visual field in V1 is largely ‘conformal’ in that it locally preserves angles in visual space (*17, 18*).

**Figure 2:**
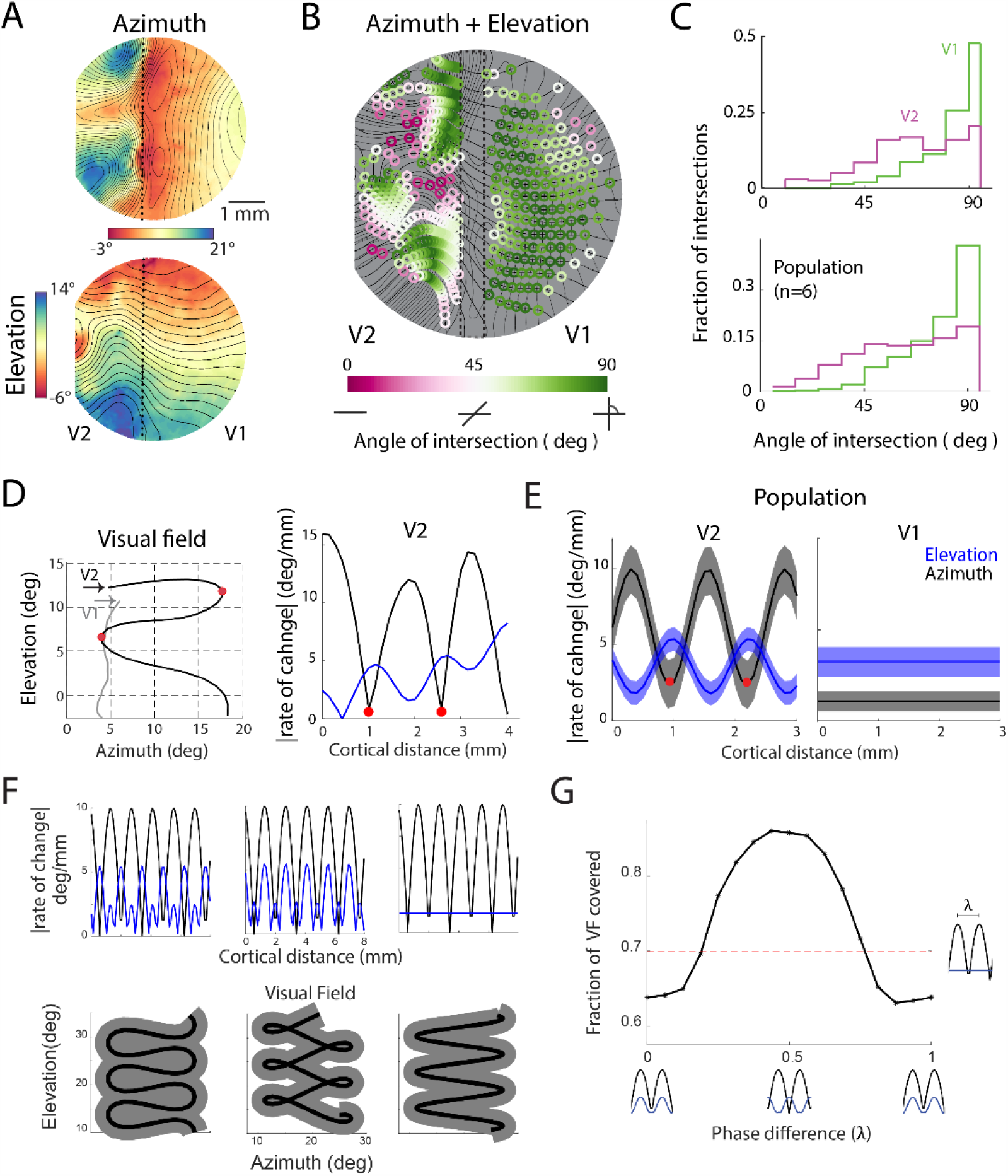
Azimuth and elevation gradients in V2 are not strictly orthogonal, and oscillate out of phase. **A)** Azimuth and elevation maps, overlaid with 1 deg contour lines calculated from maps smoothed with a 500 um nearest-neighbor filter. **B)** Overlaid azimuth and elevation contours, with intersection angle in color. **C)** Distribution of intersection angles for V1 and V2 of animal in A-B (top), pooled across additional animals (bottom), p<<0.01 two-sample Wilcoxon rank sum test. **D)** Path in visual field corresponding to linear track in V2 in Figure 1D (left). Corresponding absolute value of the rate of change in V2 path, red dots are gradient reversals (right). **E)** Population average sinusoidal fits of rate of change for paths in V2 and linear fits for paths in V1. **F)** Rates of change for azimuth and elevation with varying phase relationships including sinusoids out of phase (left) sinusoids in phase (middle) and sinusoidal azimuth but constant elevation (left), and corresponding paths in the visual field. Gray area represents receptive field size. **G)** Fraction of visual field covered vs. azimuth-elevation phase offset. The population average phase-offset (0.49) is near optimal for maximum visual field coverage.

In contrast, the oscillating pattern of the map of azimuth in V2 is accompanied by a significant departure from this conformal structure with fewer intersections exhibiting an orthogonal angle (Figure 2B, C). Thus, the orderly topological representation of visual space that has been thought to be the hallmark of visual cortical areas is not preserved in V2.

Fully capturing the spatial relationship between maps of elevation and azimuth requires considering not only the angle of the contours, but the rate of change of the contours across the cortical surface; i.e. the magnitude of the gradients (deg/mm) in azimuth and elevation and how they relate. To characterize the relative magnitude of the azimuth and elevation gradients, we consider a linear path in V1 and V2 (shown in Figure 1D) and examine the corresponding trajectory in the visual field, as well as its first derivative (Figure 2D left and right respectively). The visual field trajectory corresponding to the straight path in V1 is a line, indicating a linear transform, whereas the trajectory for V2 resembles a sinusoid (<u>Movie S1</u>). As might be expected, the gradient magnitude for azimuth exhibits a strong oscillating pattern with low magnitude at the center of the central and peripheral stripes where the gradient reverses (red dots) and high magnitude midway between the stripes. Interestingly, although the differences in gradient magnitudes for elevation are less, the elevation gradient magnitude also exhibits an oscillatory pattern that appears related to the azimuth gradients: The highest magnitude elevation gradients occur where the magnitude of the azimuth gradient is low (Figure 2D right) and vice-versa.

The inverse relationship of the azimuth and elevation gradient magnitudes is captured by fitting each gradient to a sine function (Figure S2D), and comparing the period and phase of each gradient. The period for azimuth and elevation gradients appears to be similar, 1.25+/-0.35 mm for azimuth and 1.37+/-0.36 for elevation (n=6). Two sinusoids, with similar periods, that are inversely correlated will be offset in phase by ½ the period (ƛ). For the animal shown in Figure 2D, the phase offset in V2 is 0.46ƛ. The population (n=6) average is 0.49+/-0.23 ƛ (Figure 2E left). In contrast, V1 displays a linear gradient for both azimuth and elevation (Figure 2E right). Note, in this figure the average amplitude of the gradient for elevation is higher than that for azimuth in V1 simply because the path in V1 runs perpendicular to the elevation contours.

Why should there be such a complex pattern to the map of visual space in V2? One obvious possibility is that this sinusoidal structure is related to the markedly elongated shape of V2 (*19, 20*). Unlike the relatively isotropic V1, V2 is a narrow strip of cortex that extends parallel to the border of V1 for 10-12 mm and orthogonal to the border for 1.5 mm (*19*). We reasoned that the sinusoidal transform could serve to achieve coverage of the two dimensions of visual space in a cortical area that exhibits a pronounced spatial asymmetry; a solution that effectively uses the long cortical dimension to encode changes in both azimuth and elevation (<u>Movie S2</u>). Further support for this idea comes from modeling the oscillatory patterns and phase relationships of azimuth and elevation in V2. The similar periodicity and inverse correlation is ideal for optimizing coverage of the visual field (Figure 2F-G). Indeed, the sinusoidal path we observe (Figure 2D) is reminiscent of the space-filling Peano curve in geometrical analysis and resembles the ‘boustrophedon’ pattern that has been shown to achieve optimal spatial coverage with minimal path length in the robotics path-planning literature (*21*).

Thus, the retinotopic maps in both V1 and V2 appear to be organized in a fashion that optimizes coverage of the visual field, with differences in topology necessary to accommodate differences in the shape of the two areas. A strong test of this hypothesis is to ask whether the wiring minimization principles (*22*) that have been shown to account for optimal coverage with a linear transform in V1, derive optimal coverage with a sinusoidal transform in an area with the elongated shape of V2. To address this issue, we employed the well-established elastic-net model (*3, 5*). V2 is modeled as a sheet with point ‘neurons’, each with a receptive field (RF) in the visual field (VF). Retinotopic maps are produced by color-coding V2 by the RF azimuth or elevation value of each neuron (Figure 3A, ‘Cortical Maps’). Initially, there is a random mapping between RFs in the visual field and neurons on the cortex (Figure 3A, ‘left’), although different initial conditions produce qualitatively similar results (Figure S3A). The objective of the algorithm is to arrange the RFs so they match a set of input RFs that uniformly tile the visual field (‘Target VF’), with the wiring-length constraint of preserving map smoothness – e.g. nearby neurons on the cortex should have nearby RFs in the VF. The algorithm iteratively optimizes the RFs in order to reduce the total cost associated with visual field coverage and smoothness until a stable state is reached (Figure 3A, ‘right’).

**Figure 3:**
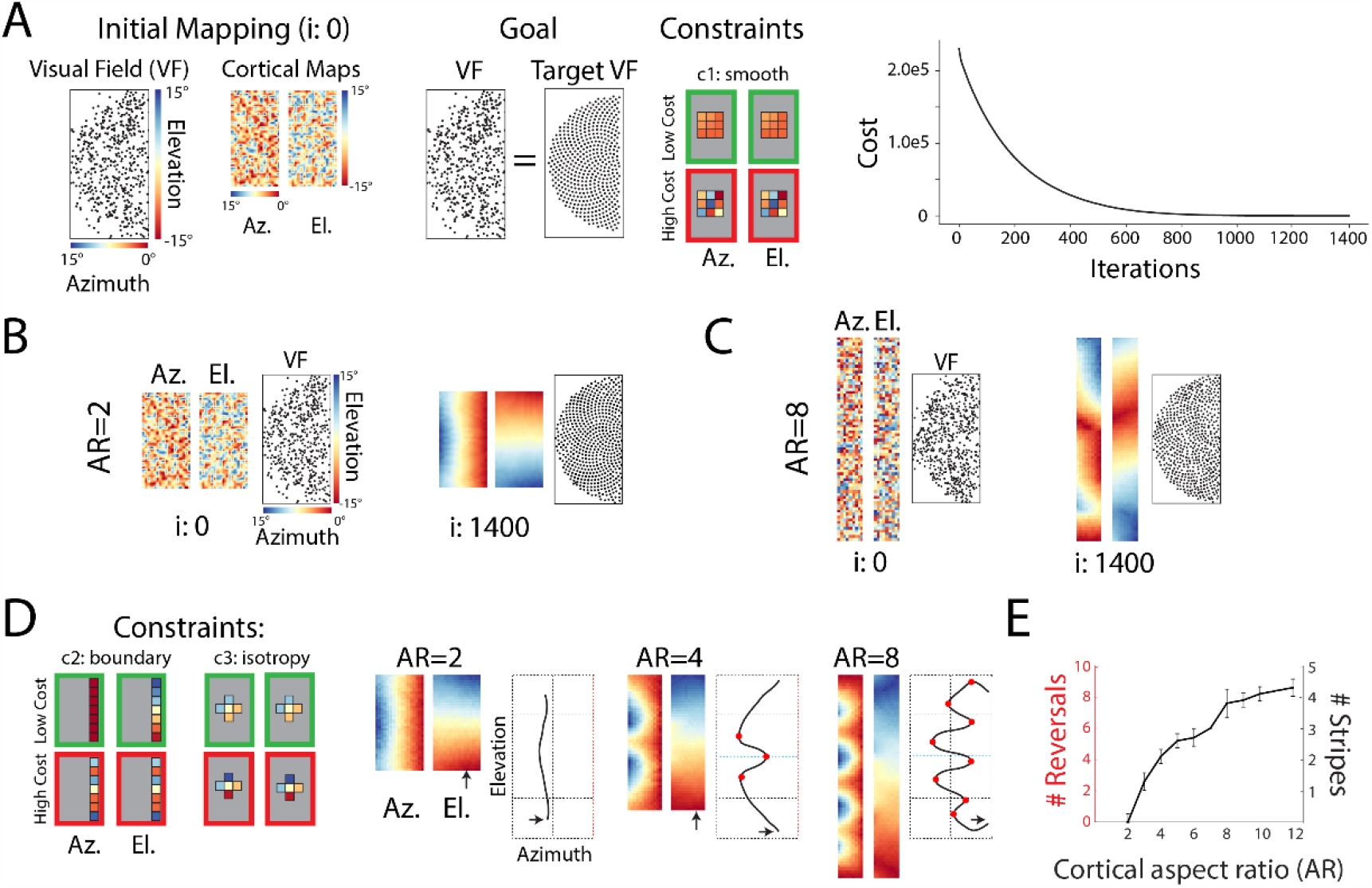
Sinusoidal retinotopy develops in elongated cortical areas. **A)** Elastic-net algorithm used to simulate the development of retinotopic maps from random initial conditions. The objective is to match RFs to an input set tiling the visual field while maintaining smooth nearest neighbor relations on the cortex (Methods). The algorithm finds solutions that lower the total cost, eventually converging to a stable solution. **B)** At small aspect ratios (AR), the final mapping consists of orthogonal maps of azimuth and elevation. **C)** At larger ARs, the final maps for azimuth and elevation are both striped along the long-axis of the cortex. **D)** Adding two additional constraints to the traditional elastic-net produces maps similar to measured data (Methods). Increasing AR produces increasingly striped azimuth maps but maintains smooth elevation maps. Traces correspond to path in VF produced by a linear track in cortex at arrowhead. Red dots are azimuth reversals. **E)** Number of azimuth reversals, and number of stripes, vs. AR. Error bars are SD, from n=10 simulations with different initial conditions.

To understand the effect of cortical geometry on the formation of maps (*20*), we varied the elongation of the cortical sheet while maintaining the total cortical area, number of neurons, and all other parameters (Methods). At small aspect ratios (AR), the optimal final mapping are simple continuous representations of the visual field, with orthogonal azimuth and elevation maps (Figure 3B). At larger ARs (Figure 3C), optimal visual field coverage is achieved by striped azimuth and elevation maps. This occurs due to the asymmetry in the number of neighbors along the two dimensions of the elongated cortex, which produces RF tiling with a one-dimensional bias (Figure S3B) that manifests as striping of the cortical maps (Figure 3C). Although the elastic-net applied to elongated cortical areas (Figure 3C) recapitulates the fundamental striped co-axial feature of retinotopic maps in V2 (Figure 1), its prediction differs from experimental data in significant ways. To improve the model we added two additional biologically inspired constraints (Figure 3D), representation of the vertical meridian at the area’s border (c2) and similarity in the local resolution along the two orthogonal axes of the cortical sheet (c3). The last constraint ensures that single RFs are not stretched significantly along horizontal or vertical axes (Figure S3D).

This full model could produce both types of maps observed in relatively isotropic V1 and anisotropic V2. As aspect ratio (AR) increased, the azimuth map became sinusoidal (Figure 3D, right) whereas the elevation map remained smooth due to the influence of the vertical meridian constraint. The oscillations that occurred in the azimuth map increased with increasing AR (Figure 3D-E), due to the local isotropy constraint, and were not dependent on the particular shape of the input RFs (Figure S3C). Similar to our experimental observations (Figure 2D), the path in the visual field corresponding to a linear track along the model cortex included several reversals (Figure 3D, red dots), which increased in number with increasing AR (Figure 3E, red). Our results confirm that the same biological principles can produce optimal coverage with linear transforms along two dimensions in isotropic cortical areas such as V1 but produce a largely one-dimensional sinusoidal transform in anisotropic areas such as V2. This is consistent with prior work on the influence of area shape and borders on retinotopic mapping (*5, 20*).

The functional data and model simulations provide evidence for a novel sinusoidal transformation of the retinal surface in V2 that is different from the simple transform in V1 (Figure 4A). The most dramatic difference between the two representations can be seen in the map of azimuth (Figure 4A, right), with the single gradient map in V1 converting to a striped map in V2 with numerous gradient reversals at the most central and peripheral regions of the visual field. Since V2 receives dense projections from V1 (*23*), we next examined whether the axonal connections from V1 to V2 are organized in a fashion that could support conversion from the map in V1 to that in V2. We began by examining the pattern of labeled axon terminals in V2 following injections of anterograde tracers in V1 parallel to, but displaced from, the V1/V2 border, thereby targeting neurons with receptive fields along an iso-azimuth axis of visual space but sparing neurons representing more central and peripheral regions. Consistent with prior observations (*24*), we found discrete stripe-like patterns of labeled terminals along the length of V2 (Figure 4B, left), as expected from the predictions of a sinusoidal transform (Figure 4B, right and <u>Movie S3</u>). This experiment is analogous to activating V1 with a vertical bar in an iso-azimuth region of the visual field (Figure 1C).

**Figure 4:**
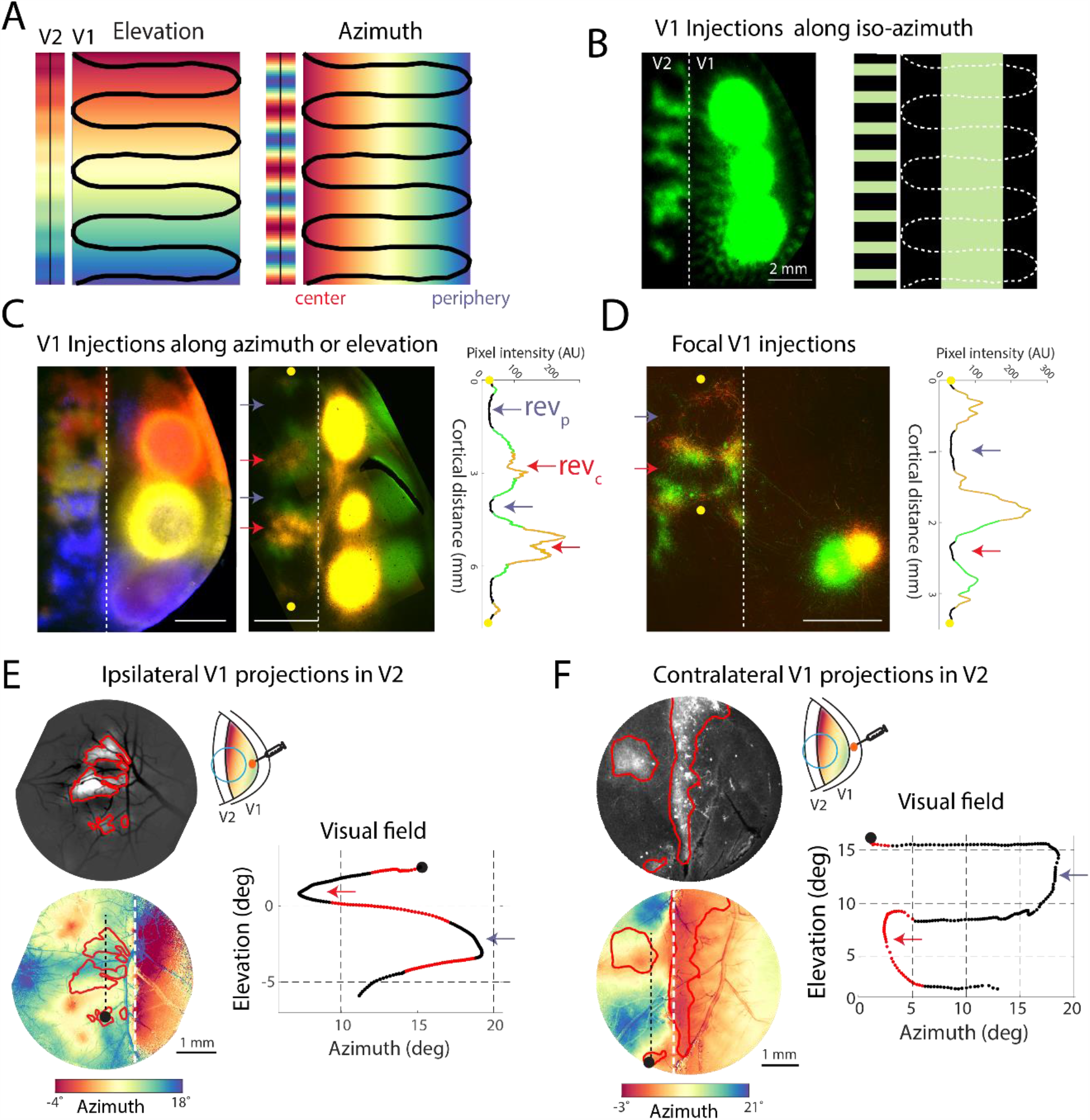
V1-V2 connectivity is sufficient to produce sinusoidal retinotopy in V2 from linear retinotopy in V1. **A)** Schematic of sinusoidal transform between V1 and V2 maps. See alternative linear transform in Figure S4A. **B)** Axon terminals in V2 produced from tracer injections in V1 along an iso-azimuth direction in flattened section of left hemisphere (left), compared to prediction from a sinusoidal transform of V1 (right). **C)** Axon terminals in V2 produced from injections in V1 along the elevation-encoding dimension (left), or orthogonal azimuth-encoding direction (right). Trace of fluorophore color and intensity values for a line in V2 shows evidence of reversals in regions representing peripheral (rev_p_) and central visual field (rev_c_). **D)** Axon terminals in V2 produced by two focal injections in V1 away from the V1/V2 border. Reversals are expected in unlabeled regions surrounded by the same color terminals, as shown in trace. **E)** Multiple axon patches in V2 produced by injection of Tdtomato in V1 outside cranial window, imaged in vivo (left). Azimuth map in same animal overlapped with axon projection contours (right). Trace shows path in visual field for the linear track in V2, with regions that pass through Tdtomato contours in red. **F)** Callosal projections produced by Tdtomato injections in contralateral V1, from flattened section (left). Azimuth map in same animal overlapped with axon projection contours (right). Trace as in F.

The conversion from the 2D linear map in V1 to a co-axial sinusoidal map in V2 also predicts that injections displaced along the elevation axis of V1 and those displaced along the orthogonal azimuth axis should both result in V2 axon terminal fields that are displaced along the same elongated dimension of V2 (Figure 4A). To test this prediction, we use different colored tracers to create coarse color maps of elevation (Figure 4C, left) and azimuth (Figure 4C, right) along orthogonal dimensions of V1. We observe that for both injection patterns, color variations in labeled terminals in V2 occur largely in the elongated dimension parallel to the V1/V2 border, consistent with our functional data (Figure 2B-C).

The pattern of labeled terminals in V2 following injections displaced along the azimuth axis of V1 is also consistent with smooth reversals in visual space evident in the functional map of V2 (Figure 1D, 4A). The axon terminals in V2 from the more central injection sites in V1 (yellow) are surrounded by the terminals derived from the more peripheral sites (green). Indeed, the spatial distribution of labeled terminals in V2 can be used to infer the location of central and peripheral gradient reversals in the map of visual space (Figure 4C, trace): Reversals in central visual field (rev_c_) would be expected in the center of the bands of yellow terminals that are surrounded by green terminals, while reversals in the periphery (rev_p_) would occur at the center of unlabeled regions surrounded by green terminals. The precision of this remapping to match the multiple reversals in V2, is also evident in patterns of labeled terminals from small neighboring injections of green and yellow tracers in V1 (Figure 4D). These injections are far from the central visual field representation near the V1/V2 border, and from the peripheral representation near the medial sinus. Each injection results in sub-millimeter thin stripes of labeled terminals in V2 that surround terminal free regions, and the stripe patterns change smoothly from green to yellow in between, reflecting the continuity of retinotopy at the two injection sites in V1.

What emerges from these experiments is a consistent pattern, where sites in V1 that represent intermediate azimuth values give rise to labeled terminals in V2 surrounding label-free zones likely to represent central and peripheral reversals in azimuth (Figure 4C-D, traces). As a direct test of this hypothesis, we combined fluorescent tracer labeling produced by a single injection in V1 (Figure S4C) with functional retinotopic maps in the same animal (Figure 4E). Labeled terminals in V2 overlapped with regions of the azimuth map representing intermediate azimuth values. By comparing the distribution of labeled terminals with the corresponding map of visual space, it is apparent that the two prominent gaps in the distribution of labeled terminals correspond to reversals in azimuth, one central, and the other peripheral (Figure 4E, trace).

We further tested the consistency between anatomical projections and retinotopic maps by injecting Tdtomato in contralateral V1 and studying the pattern of inter-hemispheric callosal terminals in V1 and V2 (Figure 4F). Callosal projections are known to terminate in regions of V1 near the representation of the vertical meridian (*25*–*30*). Our results show that this rule also generalizes beyond V1, as labeled terminals overlap with central regions in both V1 and V2 (Figure S4D). Relatively more peripheral portions of the visual field are acallosal in V2, as they are in V1, thus explaining the prominent striped pattern of callosal projections previously observed in tree shrew V2 (*25*) (Figure S4E). These results demonstrate that the connections from V1 to V2 exhibit the fine-scale topography that is sufficient to build the sinusoidal co-axial transform in V2 from the linear orthogonal transform in V1.

Does the sinusoidal retinotopy have significance for understanding the spatial distribution of functional properties in V2? Given the juxtaposition of relatively more central and peripheral regions of visual space and the selective targeting of callosal inputs to the central regions we wondered whether this would be accompanied by differences in the processing of binocular information. Similar relationships between degree of binocularity and visual field representation have been reported in V1, where neurons in the representation of the vertical meridian are more sensitive to binocular stimuli (*31*). To determine whether V2 exhibits a map of relative binocularity we measured the response to retinotopy stimuli viewed binocularly, or monocularly through the contralateral eye, and subtracted them to highlight relative differences in binocularity (Methods). We found interleaved regions of V2 that were relatively more binocular or monocular, and the overall pattern was highly correlated to the azimuth map in the same animal (Figure 5A). As predicted, V2 regions that represented more central regions of the visual field were more binocular compared to regions that represented more peripheral regions.

**Figure 5:**
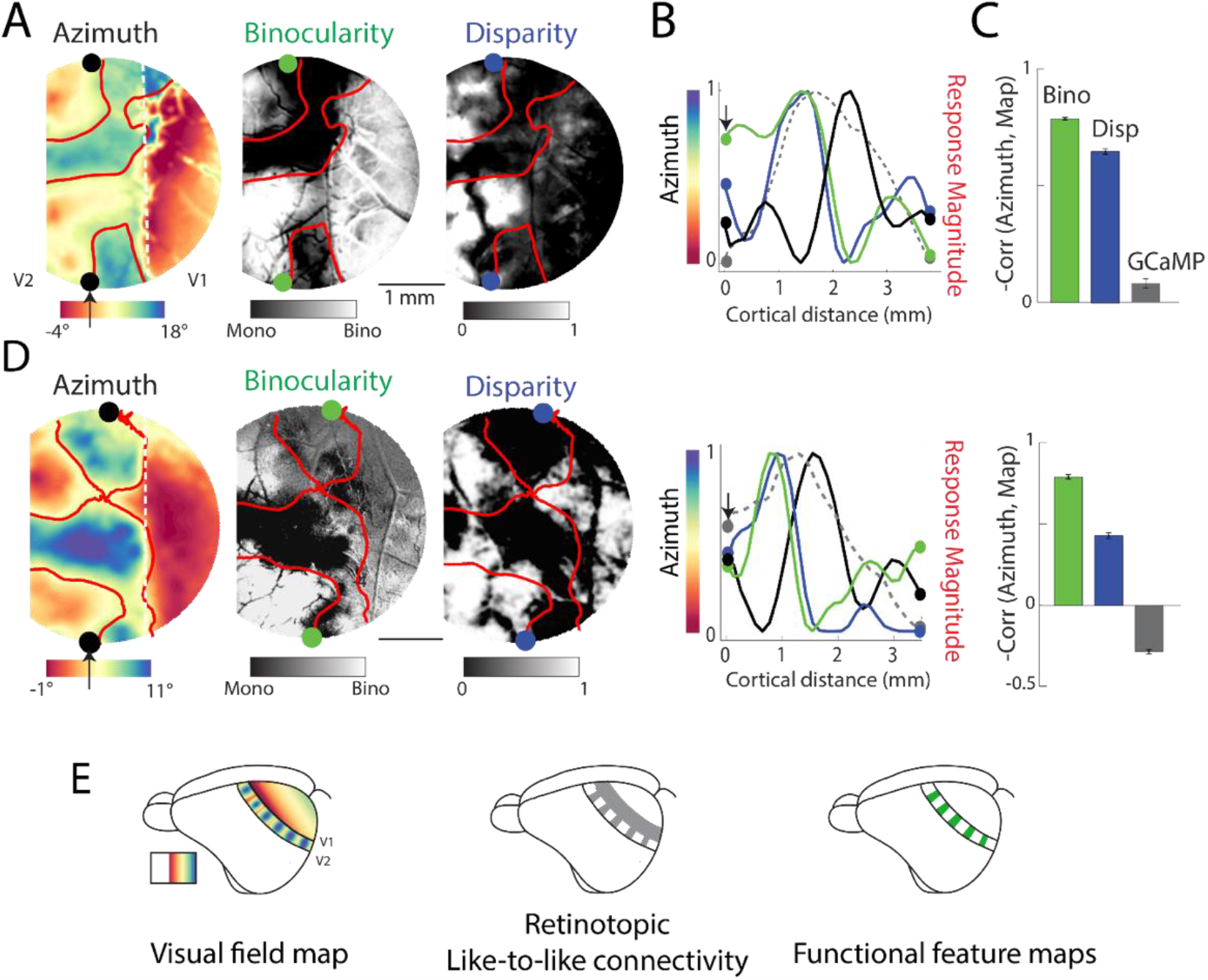
Functional feature maps in V2 are correlated with the azimuth map. **A)** Functional maps in the same animal. Azimuth map overlaid with contour line indicating mid-point of azimuth. Map of relative binocularity and relative horizontal disparity sensitivity (Methods). All feature maps are thresholded ±1 SD of the mean and normalized. **B)** Extracted pixel values for the azimuth and normalized feature maps corresponding to a line in V2 indicated by arrowhead in A. **C)** Mean pixel-wise correlations between the azimuth and feature maps in V2, Bino = −0.78±0.01, Disp = −0.71±0.01, GCaMP -0.08±0.02. Negated correlation values are shown for easier interpretability. Error bars are SEM of 1000 bootstraps. **D)** Same as A for a different animal. Bino = −0.79±0.01, Disp = −0.44±0.02, GCaMP 0.28±0.01. **E)** Visual field map can explain retinotopic like-to-like connectivity between V1 and V2 and within V2 and correlated striped appearance of functional feature maps.

Given such differences in binocularity between relatively central and peripheral stripes, led us to wonder whether binocular disparity - a fundamental cue for depth information - would also be differentially represented across V2 stripes. Using two-photon single-cell imaging, we confirmed that neurons in V2 are disparity tuned (Figure S5A). For widefield imaging we were interested in measuring the relative disparity sensitivity across the stripes of V2. We therefore measured the pooled response to red/green random-dot stereograms drifting in various directions with and without horizontal disparity cues and subtracted the two resulting maps (Methods). As might be expected, stripes of V2 that represent more central regions of visual space exhibited higher sensitivity to horizontal disparity than those representing more peripheral regions. (Figure 5A).

We thus found that the striped azimuth map in V2 was correlated with functional feature maps. We extracted the pixel values along a line in V2 running across the stripes, at the same location in all maps (Figure 5B). The co-localized maxima and minima of the normalized curves corresponding to feature maps (colored lines) indicates their similarity along a line in V2. We calculated the pixel-wise correlation between the azimuth map and feature maps (Figure 5C) and found the strongest correlation between the azimuth and binocularity map (0.79 ± 0.01 SEM of 1000 bootstraps). Furthermore, we found small or oppositely signed correlations between the pattern of GCaMP expression and the azimuth map (−0.28± 0.01, Figure S5B), ruling out the possibility that these correlations are simply due to a shared and non-uniform fluorophore expression pattern. Data for a second animal is shown in Figure 5D. In summary (Figure 5E), we found that the oscillating pattern of visual field representation we observed in functional and anatomical data are correlated with a similar oscillating pattern of functional feature properties. Further investigation of other response properties, such as color and orientation, may reveal retinotopic anchoring in the organization of other functional feature maps.

## Discussion

From the earliest descriptions of retinotopic maps in the brain, accumulated evidence has supported a fundamental principle for retinotopic organization best described as a conformal transform. Our results challenge this perspective, demonstrating a novel sinusoidal retinotopic mapping that exhibits the fine scale properties ascribed to traditional maps--uniform coverage of visual space, adherence to wiring minimization-- and is optimal for achieving this within the elongated spatial constraints of cortical area V2.

The departure from linearity that we have described should not be confused with fovea related spatial distortions of the linear map of visual space or the fine-scale discontinuities in the mapping of visual space that have been suggested to accommodate ocular dominance columns in V1 (*32, 33*). Our observation of a single sinusoidal representation of the visual field is also inconsistent with multiple interleaved representations, as reported in primate V2 (*11, 34*). Instead we show a fundamental difference in the global structure of the retinotopic map where the conventional orthogonal relationship of azimuth and elevation is replaced with a broad range of intersection angles that allow azimuth and elevation to co-vary along the same axis in cortical space. The stripes we observe in the retinotopy map are merely reflections of the sinusoidal transform, and not indicative of multiple representations since regions that represent similar azimuths do not represent similar elevations.

These results demonstrate that the form that retinotopic maps can take is far more flexible than we appreciated, and that we have much to learn about the mechanisms that shape the topological structure of neural representations (*35, 36*). While core principles of uniform coverage and wiring minimization appear to be conserved in retinotopic maps, other traits such as lack of reversals and preservation of orthogonal axes are clearly abandoned. Our results suggest that features of the cortical area, such as its shape, boundaries with adjacent areas, and sequence of development can dramatically alter the nature of the transform and corresponding map layout (*5, 35*). Furthermore, the inherently one-dimensional nature of the sinusoidal transform we observe suggests that cortical surface representations must not be constrained to exactly two dimensions. It remains to be seen what other transforms might be found in the retinotopic maps that comprise the large number of cortical areas that extend beyond V2.

The significance of these results extends far beyond the transform itself, forcing a re-consideration of the role of retinotopy in the organization of intracortical connections, and the spatial layout of functional feature maps (*33, 37*). Indeed, stripe-like patterns of connectivity and functional properties have been described in numerous studies of tree shrew (*24, 25*) and primate V2, but these observations have been interpreted as clustering of functionally distinct streams that are independent of the map of visual space (*11*). Likewise, the striped organization of callosal connections in V2 has been recognized in macaques (*38, 39*) and other species (*25*–*27, 40*) as potentially reflecting a bias of these connections for central regions of visual space, but their relationship to visuotopy has remained enigmatic (*28, 39*). Given the elongated shape of area V2 in most species (*41*), the sinusoidal retinotopic organization we have described in tree shrew is likely to provide an explanation for the stripe-like arrangement of response properties and connections that is evolutionarily conserved. Notably, close inspection of receptive field positions measured with electrode tracks parallel to the V1/V2 border in Cebus monkey (*42*), macaque (*11, 34*), and marmoset V2 (*43*) reveals small oscillations in the representation of the visual field similar to the tree shrew (Figure 1).

Implicit in the correlation of sinusoidal retinotopy with differences in functional properties and connectivity is the presence of functional specializations within V2 that are specific to different regions of visual space. Some of these specializations such as binocular responsiveness and tuning for disparity are not surprising given the relatively modest amount of binocular overlap (30 degrees) in the field of view of the two eyes of the shrew (*16*). Nonetheless, the sinusoidal organization of V2 spotlights functional specializations within the retinotopic map that are likely to reflect distinct and behaviorally relevant processing of information from different regions of visual space. This is broadly consistent with different variations of the ‘two streams hypothesis’ (*44*–*46*). As such, we predict that comparative analyses of the functional properties that exhibit clustering within macaque V2 (e.g., color, direction, etc.) will reveal additional species-specific visual field specializations that have evolved to support niche-adaptive visual behaviors.

Lastly, our work highlights the powerful role of retinotopy in anchoring the anatomical and functional organization of cortical circuits within the visual hierarchy (*47*). Understanding the transform between retinotopic maps in areas along the hierarchy can produce precise predictions of inter and intra-area connectivity and interhemispheric projections. Thus, high-resolution whole brain imaging approaches can largely constrain the wiring diagram of the visual cortical hierarchy, allowing more efficient anatomical reconstruction, ultimately leading to better understanding of visual perception.

## Acknowledgements

We thank Susan Freling, Solana Liu, and Jessica Kerr, as well as the Animal Resource Center at the Max Planck Florida Institute for Neuroscience for help and expertise with animal care and training. We also thank members of the Fitzpatrick lab, as well as Seneca Harberger and Matthew Krause for comments and discussion. This work was supported by R01 EY006821 (DF), F32 EY026463 (MSS), and the Max Planck Florida Institute for Neuroscience.

## Materials and Methods

All experimental procedures were approved by the Max Planck Florida Institute for Neuroscience Animal Care and Use Committee and performed in compliance with guidelines published by the National Institutes of Health. Tree shrews (*Tupaia belangeri*, n = 13, 8 months-2 years old, 140-210 grams, male and female) were used for chronic recordings while awake and/or for tracer injections followed by histology. Animals were housed in single occupancy cages in a 12 hour lights on/off animal suite. Their food and water aliquots were given once daily but both were enough to maintain their weight and not considered as ‘restricted’. Animals intended for chronic recording went through a 1-3 week period of behavioral acclimation to the awake recording setup. Some animals were water restricted during this period to help acclimation, although even the minimum daily water amount (15 mls) was enough to maintain their baseline weight. Following this period, a sterile surgery was performed to inject genetically encoded calcium indicators into the visual cortex. Following an expression period of 2-4 weeks, awake animals were imaged during several experiments, each lasting 0.5 to 1.5 hours. In rare cases when imaging sessions occurred after several weeks ‘off’, animals were given low doses (0.2x dose below) of a sedative (Midazolam) to reduce stress during imaging sessions.

### Craniotomy and intra-cortical expression of viral vectors

Tree shrews were sedated with Midazolam (5 mg/kg, intramuscular (IM)), anesthetized with ketamine (75 mg/kg, IM) and given atropine (0.5 mg/kg, subcutaneous (SC)) to reduce secretions. A long-acting analgesic (slow-release Buprenorphine, 0.6 mg/kg, SC), as well as dexamethasone (1 mg/kg, IM) to reduce swelling, was administered before the surgery. The animal’s head was shaved, cleaned, and the surgical site was injected with a mixture of bupivacaine and lidocaine (0.3–0.5 ml, SC) to reduce local pain. Isoflurane anesthesia (0.5–2.5%) was initially delivered through a nose-cone, and once the animal was stably anesthetized, via an endotracheal tube. A mechanical ventilator provided respiration at 100 strokes per minute. The animal’s EKG was monitored through leads placed on the chest and back. Venous cannulation (hind limb) was performed to deliver 5% dextrose solution. Internal temperature (36–37 °C) was maintained by a thermostatically controlled heating pad while expired CO_2_ and heart rate were monitored to assess anesthetic depth and overall health. The animal was placed in a stereotaxic device (Kopf, Model 900), a small incision was made over the midline, skin and muscle were retracted. A metal headplate was cemented (Metabond) to the anterior skull over the olfactory bulbs and a metal chamber (inner diameter 8 mm) was cemented to the skull over the visual cortex. A craniotomy was made within this chamber (about 6-mm diameter), centered at 4A, 4L relative to lambda, using a motorized jeweler’s drill. The visual cortex was injected with a total of 2–3 μl of virus solution (1e13 GC/ml) containing AAV9-Syn-GCaMP6s.WPRE.SV40 (Penn Vector Core or Addgene) through a beveled glass micropipette (tip size 10–20 μm diameter) using a nanoinjector (Drummond Nanoject II, WPI). A total of 6-10 injection sites were made in each cranial window, while the dura remained intact. To facilitate spreading, the virus was delivered at two depths, 250 and 500 μm below the cortical surface. After the injections, a durotomy was performed, and a 3-5 mm glass window, glued to an 8 mm coverglass was placed inside the craniotomy and sealed with Vetbond. The surface of the metal chamber was covered with black dental cement to prevent light reflection. The skin around the incision was sutured closed. The animals were then placed on a heating blanket in a small cage to recover from anesthesia. The first imaging experiments occurred following a period of 2-3 weeks. The animals continued their acclimation to the recording apparatus during this time. Animals were given twice daily doses of Baytril (5 mg/kg) for 5 days to reduce the probability of infections.

### Chronic recording setup and head fixation

All data, except that in Figure S3D were recorded in awake animals passively viewing visual stimuli on a monitor placed parallel to the eyes. The data in Figure S3D was recorded in isoflurane (1%) anesthetized animals. Animals were head-fixed to a home-built stereotax using metal screws that interfaced with the metal head-plate previously cemented to the skull. The stereotax also included a clear acrylic tube that served to constrain the animal’s body movements. Animals were acclimated to this setup prior to and after the injection surgery using juice rewards. During imaging experiments, some animals were rewarded with juice after the completion of stimulus blocks, but no animals received juice within stimulus experiments.

### Calcium imaging experiments

Wide-field 1-photon and two-photon calcium imaging was performed using the Bergamo II Scope (Thorlabs) with blue LED illumination using an LED driver (Thorlabs). GCaMP6s fluorescence signal from the cortical surface was acquired at about 15 Hz (640 × 540 pixels, field of view (FOV) ranges from 3×3 to 5×5 mm^2^) using a Zyla 5.5 sCMOS camera (Andor) controlled by μManager2. Average excitation power at the exit of the air objective (2x or 4x, UPlanFl, Olympus) ranged from 0.2 to 0.8 mW. Individual sessions were initially registered prior to recording using a blood vessel template recorded on the first chronic session. More accurate registration was performed offline using custom scripts based on pixel correlation. Single-condition response maps were expressed as 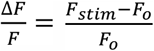 where *F*_*stim*_ is the trial and time-averaged response across stimulus frames, and *F*_*o*_ is the trial and time-averaged response during the 1 second prior to stimulus onset.

### Two-photon imaging experiments

Imaging experiments were performed using the Bergamo II Scope (Thorlabs) with either 920 nm excitation provided by an InSight Dual (Spectra-Physics) or 920 nm excitation provided by a Mai Tai DeepSea laser (Spectra-Physics), controlled by ScanImage software (Vidrio Technologies). Average excitation power at the exit of the objective (16×, CFI75, Nikon Instruments) ranged from 20 to 50 mW. Images were acquired at 15 Hz (512 × 512 pixels, field of view (FOV) ranges from 0.44 × 0.44 mm^2^ to 1.1 × 1.1 mm^2^). Imaging depth was between 75 and 200 um below the surface, corresponding to Layer 2/3 of the tree shrew visual cortex. Z-projections of two-photon fields of view were aligned to the epi-fluorescence imaging with the blood vessel pattern. Registration and Single-cell ROIs selection was done automatically using Suite2P ((*48*) https://github.com/MouseLand/suite2p). The response to each stimulus was calculated as 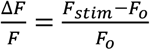 where *F*_*stim*_ is the response during each frame when the stimulus was on, and *F*_*o*_ is the time-averaged response during the 1 second prior to stimulus onset. Except where noted, visual stimuli for 2-photon experiments were presented for 3 seconds, with a 3 second interstimulus interval.

### Visual stimulus methods

Visual stimuli were delivered on a monitor using PsychoPy (*49*). The monitor had resolution 1920×1080 pixels and at the typical distance of 20 cm from the eyes extended ∼120 x 70 degrees. The center of the monitor, defined as (0,0) was matched to the mid-point between the two eyes. As described above, the animals were stationary and head-fixed during passive viewing of all visual stimuli, except the coarse receptive field mapping stimuli in Figure S3D. The eyes were on a horizontal plane (the microscope rotated to achieve parallel plane). The monitor was placed parallel to the eyes at a distance of 20 cm.

Retinotopic mapping stimuli consisted of moving black bars on a gray screen, with a width of 3 degrees, which moved at 4 degrees per second either horizontally for phase-encoded azimuth mapping or vertically for phase-encoded elevation mapping. Between 30-50 trials were used for each condition. For each new animal, the first stage of retinotopy mapping included showing full-screen stimuli to discover the limited range of the visual field that activated neurons in portions of V1 and V2 within the cranial window. This was done with the monitor parallel to the animal (Figure S1A) at a distance of 20 cm, covering 116 degrees of azimuth and with the monitor placed at 45 degrees relative to midline, approximately parallel to the contralateral eye at a distance of 13 cm, to cover up to 130 degrees of azimuth space. For monitors placed close (13 cm) to the eye, spherically corrected stimuli were used (*50, 51*). Following this procedure, stimuli were then shown in the relevant limited portion of the visual field.

To determine the relative sensitivity to monocular or binocular visual stimuli, moving edge stimuli were shown in the binocular condition, with both eyes un-occluded, and in the monocular condition, with the contralateral eye occluded. The moving edge alternated between a dark edge moving against a white background and a white edge moving against a dark background, covering 25 degrees of the ipsilateral and contralateral visual field. Only vertical moving edges were used and 30-50 trials were run. Binocular sensitivity maps produced using drifting gratings produced qualitatively similar results. All other stimuli were shown in the binocular condition.

To evoke responses related to horizontal disparity, correlated red/green random dot stereograms were used in conjunction with red/green filtered goggles placed on the animal’s eyes. The combination of the red/green filters and the relative position of the red and green dots on the monitor produced stereograms with uncrossed and crossed disparities in a range between +1.5 and -1.5 degrees. Dots drifted in 1 of 8 different directions, and had a range of inter-ocular horizontal disparities encoded by a range of different horizontal distances between the center of red and green dots. At zero disparity, the red/green dots merged into a single yellow drifting dot. The stimulus covered ±30 degrees of azimuth and elevation. The remaining background space of the monitor was filled with static yellow dots with the same density as the moving dots within the visual field. 10 trials were run for disparity measurements at the wide-field and two-photon level. For wide-field maps exhibiting disparity sensitivity, each trial consisted of 8 directions of drifting dot motion and two disparities (0, -1.5). For two-photon measurements, the goal was to quantify the disparity tuning curve, therefor each trial consisted of a single direction, corresponding to the preferred direction of motion in the field of view, and 13 disparities. Stimulus duration was 2 seconds, with a 2 second inter stimulus interval.

To map receptive fields, the relevant portion of the visual field corresponding to the two-photon field of view was identified using small apertures with drifting gratings at all orientations. A 20×20 degree box was centered over this region, and split into 2×2 degree squares. Each square was randomly flashed for 0.5 seconds, followed by a 1 second inter-stimulus-interval. 20 -30 trials were repeated.

### Tracer injections and histology methods

Some animals were injected with viral tracers in V1, in order to map out the rules of connectivity between V1 and V2. Animals in which we recorded functional calcium data were also processed for histology. Viral tracers included AAV9-Syn-GCaMP6s.WPRE.SV40, AAV1.CAG.tdtomato.WPRE.SV40, AAV1.CAG.Ruby2sm-Flag.WPRE.SV40 and AAV1.CAG.GFPsm-myc.WPRE.SV40. The GCaMP6s virus was used for green fluorescent tracer labeling, the Tdtomato for red, and the two spaghetti monster (*52*) variants were paired with secondary fluorophores in either blue or far-red. This allowed us to get up to 4 color labeling in one animal. Viral tracers were injected intracranially as described in the surgical procedures above. Perfusion and histology was done following a 3-4 week expression period.

Animals received a lethal injection of the barbiturate Euthasol and were perfused transcardially with 0.9% NaCl in 0.1 M PB, followed by 2 or 4% paraformaldehyde (PFA). Once the brain was extracted, the two hemispheres were separated and each cortical hemisphere was flattened in PFA between two glass slides with a weight on top. After 2 hours, the flattened hemispheres were submerged in PFA and stored overnight at 4 °C. After 24 hours, the hemispheres were transferred to a 30% sucrose solution in phosphate buffer and stored at 4 °C until sectioning. 50-um-thick were sectioned parallel to the flattened surface using a freezing microtome. For immunostaining, the slices were incubated in blocking solution for 30 min and then transferred to the primary antibody solution for overnight incubation at 4 °C. The slices were then incubated in secondary antibody for 2 h at room temperature, mounted on glass slides, dried, and coverslipped. Labelled neurons and structures were imaged on a confocal microscope.

### Data Analysis

Analysis was performed using custom scripts in Matlab or Python.

Azimuth and elevation, produced by moving bar stimuli (Figure 1A), were phase-encoded, where the temporal phase of the horizontal or vertical drifting bar directly encodes iso-azimuth and iso-elevation coordinates on the stimulus monitor. Therefore estimating the temporal delay of the response for each cortical pixel can be used to localize the region of the visual field to which the pixel neurons respond best. For each cortical pixel, the phase of the first harmonic at the stimulus frequency (1/period) was determined using a discrete Fourier transform of the trial-averaged time-series for each motion condition. 30-50 trials were run for each condition. The stimulus-response delay due to the rise-time of the calcium indicator was calculated by subtracting phase maps produced by bars moving in opposite directions. Delay-corrected maps were produced by subtracting this delay. Finally, maps were smoothed with a zero-phase digital ‘hamming’ filter with window width of 100 um.

For retinotopy maps, an iterative algorithm was used to produce blood-vessel free maps (compare Azimuth map in Figure 1D and Figure 4F). A highpass Gaussian spatial filter was used to detect the outline of blood vessels, producing a blood vessel outline. The mask was then filled to produce a blood vessel mask. Pixels within the blood vessel mask were removed from retinotopy map images and replaced with an average of their nearest neighbors, produced with a hamming filter of size 200 ums. The medial and lateral border of V2 were determined based on the retinotopic maps. As shown previously, the lateral portion of V1 consists a compressed representation of up to 30 degrees of the ipsilateral visual field, the lateral edge of which aligns well with the cytoarchitectonicaly defined border of V1 (*10*). We used the lateral edge of this region to designate the V1/V2 border (Figure S1B, top). In animals where we had combined functional maps and anatomical tracer data, this border coincided well with the lateral edge of callosal terminals (Figure 4F) and the medial edge of terminals produced by injections in ipsilateral V1 (Figure 4E). For demarcating the lateral border of V2, we used the midpoint of discontinuity in the elevation map (red blob in Figure S1B, bottom). This discontinuous representation of the lower visual field is related to the border of a nearby visual area TD (*19*). Further studies are needed to gain insight into the retinotopic maps of visual areas beyond V2. This demarcation was consistent with the known width of V2 based on previous cytoarchitecture studies(*19*) and our own tracer injection studies.

Maps of relative binocularity (Figure 5) were calculated from subtracting phase-encoded azimuth maps estimated during binocular viewing from azimuth maps estimated during monocular contralateral viewing, while the ipsilateral eye (left eye for a left cranial window) was physically blocked from viewing the monitor. The azimuth maps for each condition were produced from responses to identical moving edge stimuli, as described above. The monocular map was subtracted from the binocular map and the resulting difference was thresholded between mean ± 1SD.

Disparity analysis was carried out at both single-cell and wide-field scales since there has been little prior work on disparity sensitivity in the tree shrew visual system. For two-photon single-cell recordings, drifting random dot stereograms drifted at the preferred direction of the FOV and each trial consisted of 13 inter-ocular horizontal disparities ranging from -1.5 to 1.5 degrees. This allowed us to produce single-cell disparity tuning curves (Figure S5A). For widefield recordings covering regions with many different direction preferences, stereograms were shown with 8 drifting directions and only two horizontal disparities (−1.5 and 0). The single-condition maps for each direction and horizontal disparity were calculated using the 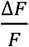 as described above. Then the single-condition maps were vector-summed (without angle doubling) for all 8 orientations to produce a single-condition-disparity map at -1.5 degrees and another at 0 degrees. To produce the disparity sensitivity map (Figure 5), the 0-degree map was subtracted from the -1.5-degree map.

All functional feature maps were thresholded between mean ± 1SD where mean and SD are calculated across all pixels, and then normalized to the unit scale to allow comparison among maps.

### Model methods

We used the elastic net algorithm (*3, 5*), and modified it for the formation of visual field maps in V2. We refer to (*5*) and our published interactive code for the implementation of this algorithm (https://github.com/msarvestani/sinusoidalTransform), and briefly describe the framework here.

V2 is modeled as a rectangular sheet, with variable aspect ratio, but fixed area consisting of 600 pixels or ‘neurons’. Each neuron has a point RF in the visual field, defined by its azimuth and elevation values. The neurons are initially arranged randomly on the visual field and cortex. The objective of the iterative algorithm is to arrange the neurons, using gradient descent, to minimize the cost function.

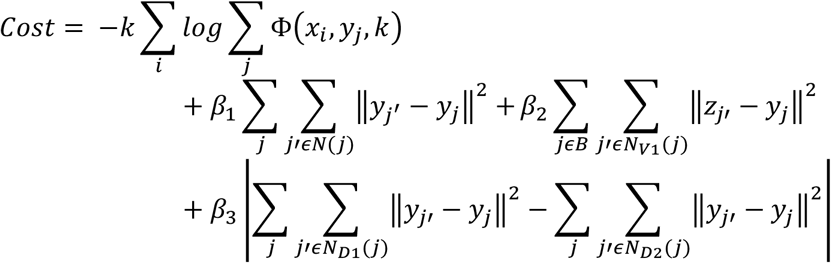

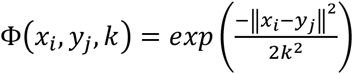, k is an annealing factor that reduces on each iteration.

In simple terms, the target RF positions (*x*_*i*_) are provided (Figure 3A, ‘Target VF’). At each iteration, the cost function produces a trade-off in minimizing the Euclidian distance between the RFs *(y*_*j*_) and the target set of RFs (*x*_*i*_) with several constraints applied to the relative position of neurons on the cortical surface. The first term is the main cost which is minimized when *y*_*j*_ is close to *x*_*i*_, assuring optimal visual field coverage as in the ‘Target VF’. The second term, weighed by *β*_*1*_ is minimized when RFs, and their nearby neighbors *N* on the cortex, are similar, ensuring smooth cortical maps. The third term, weighed by *β*_*2*_, is minimized when neurons at the border of the cortical area *N*_*V1*_have RFs close to the vertical meridian. The border that V2 shares with V1 maps the vertical meridian in tree shrews and many other species. The fourth term, weighted by *β*_*3*_, is minimized when the RFs have nearby neighbors that are similarly close along the two orthogonal directions of the cortical sheet. This ensures that the population receptive field for a group of nearby neurons on the cortical surface is relatively isotropic and not significantly stretched.

Parameters used: *k* =100 and reduces by 0.05% on each iteration. *β*_*1*_ = *β*_*2*_ *= β*_*3*_ = 0.03, except for simulations in Figure 3B-C where *β*_*2*_ *= β*_*3*_ = 0. We note that not all parameter choices produce qualitatively similar results as those shown in Figure 3. We chose parameters that produced smooth orthogonal maps at low aspect ratios, and observed changes in cortical maps that occurred as a function of increasing the aspect ratio.

## Supplementary Figures

**Figure S1:**
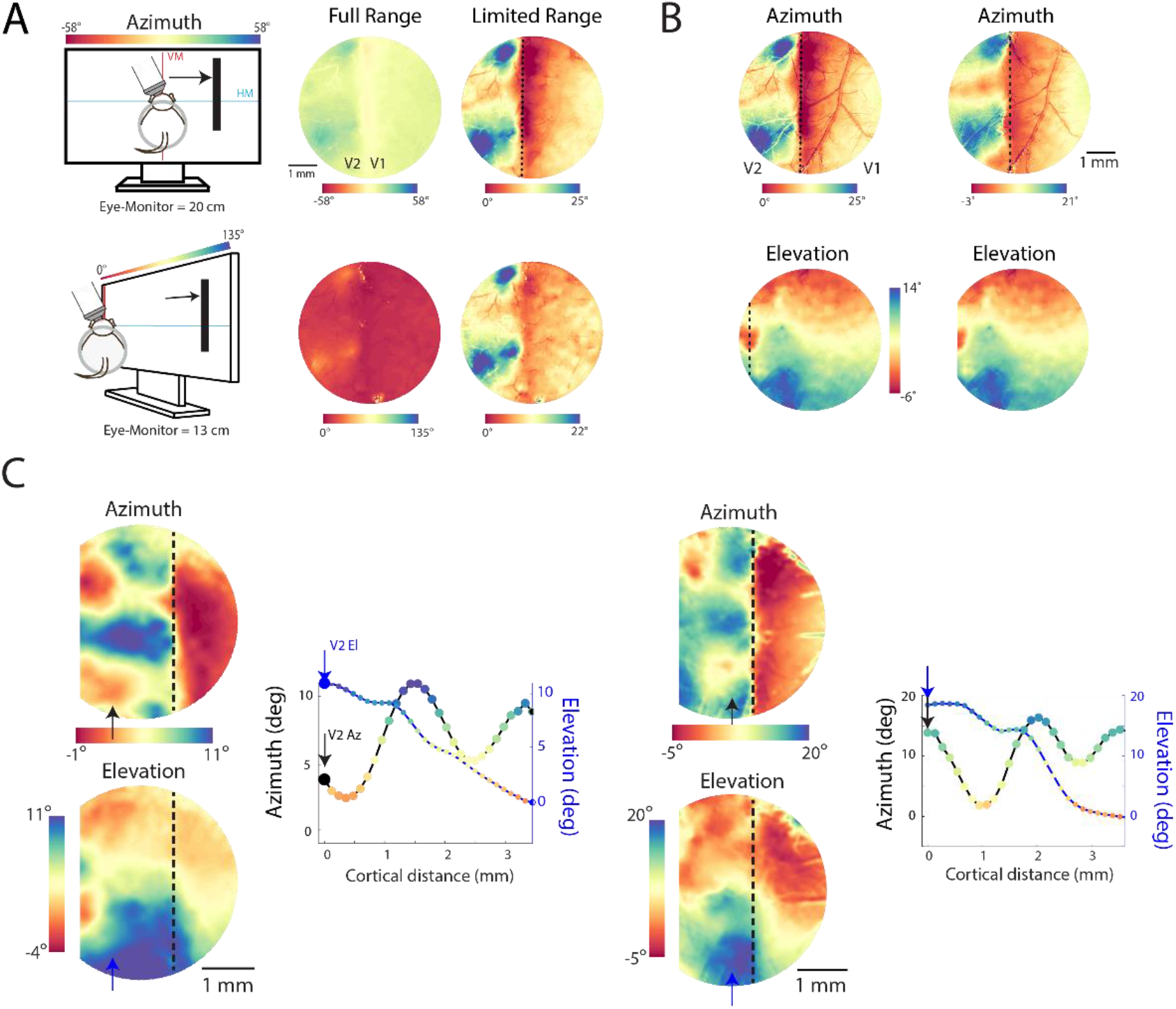
Supporting information for Figure 1. **A)** For each new animal, the first stage of retinotopy mapping included showing full-screen stimuli to discover the limited range of the visual field that activated neurons in the cranial window, with the monitor parallel to the line connecting the two eyes, as well as parallel to the contralateral eye (Methods). **B)** The V1/V2 border was demarcated using the lateral edge of the compressed ipsilateral representation in V1 (Methods). The lateral edge of V2 was determined using the elevation discontinuity related to area TD (Methods). **C)** Retinotopy maps for two additional animals. Projected azimuth and elevation values are shown for a linear track in V2. Plotting these values against each other produces the path in the visual field, with path reversals indicated with red dots.

**Figure S2:**
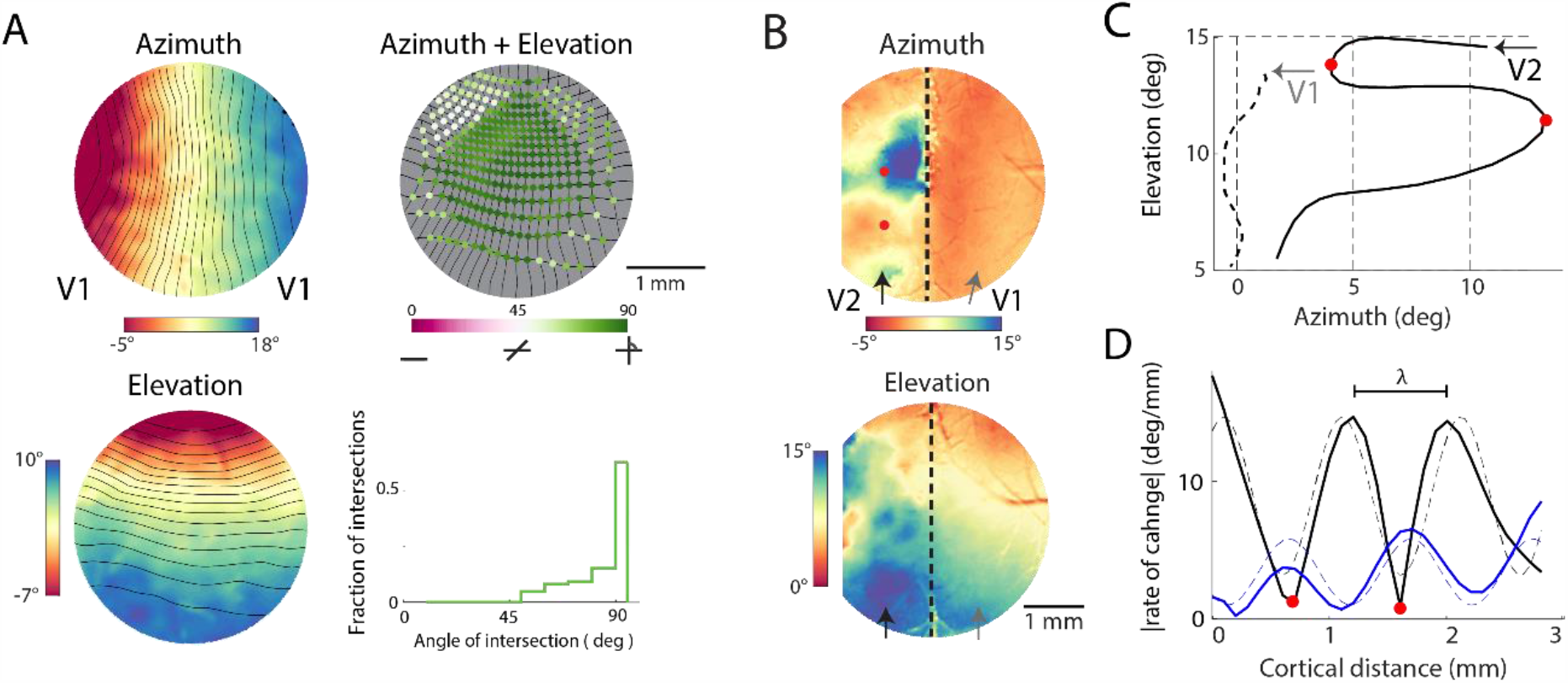
Supporting information for Figure 2. **A)** Azimuth and elevation maps in V1, overlaid with 1 degree contours calculated on maps smoothed with a 500 um nearest-neighbor filter. The 3 mm cranial window covers more lateral portions of V1 away from the V1/V2 border. The angle of intersection of azimuth and elevation contours are biased towards 90 degrees. **B)** Azimuth and elevation maps for a different animal with arrows indicating linear tracks on the cortex. **C)** Path in visual field corresponding to linear tracks in B. **D)** Absolute value of the rate of change for azimuth and elevation gradients, overlaid with fits to sinusoids with period λ. Red dots indicate gradient reversals.

**Figure S3:**
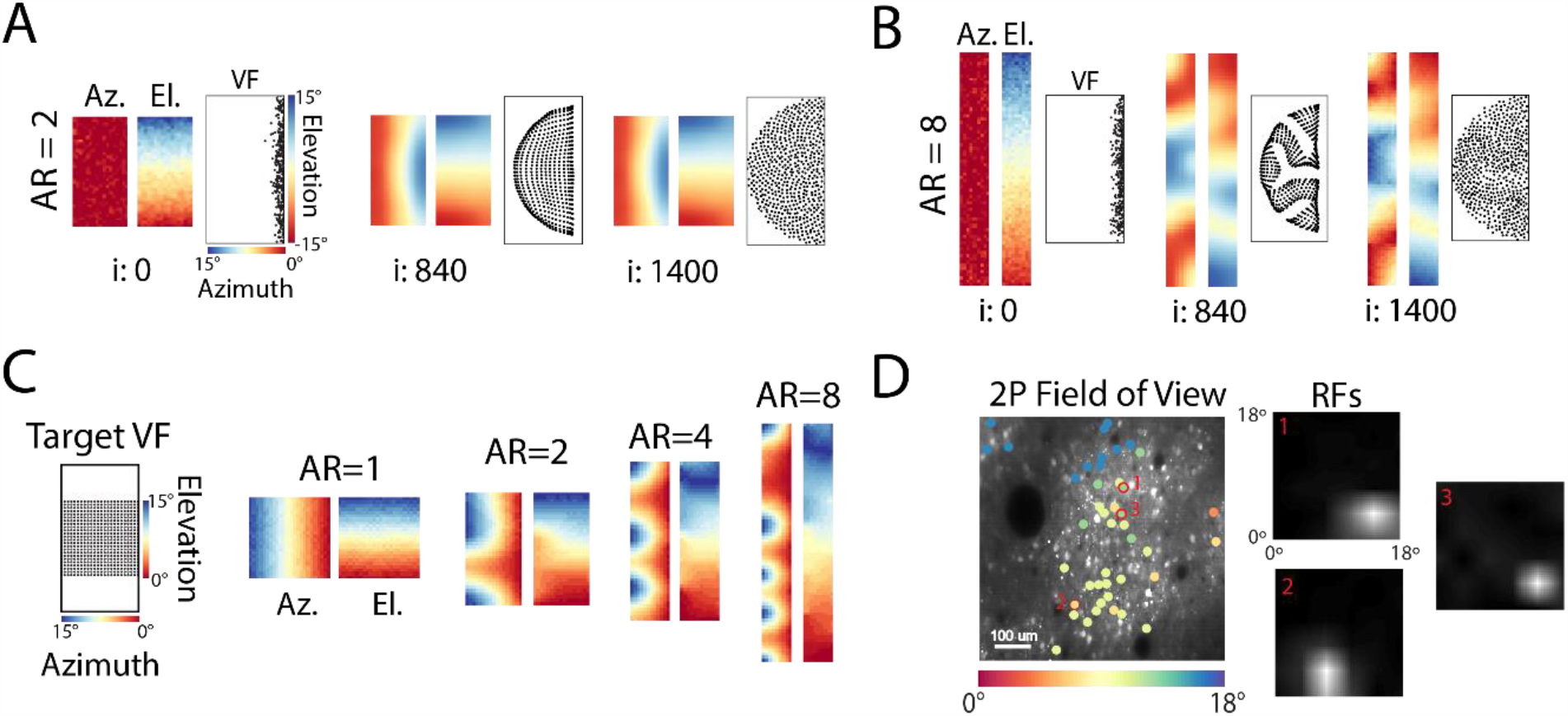
Supporting information for Figure 3. **A)** Maps produced from initial condition where all RFs lie along the vertical meridian, are similar to maps produced by initial random conditions in Figure 3B. This initial condition makes sense in light of developmental studies that show the early formation of the vertical meridian. Before convergence to stable maps (i:840), the RFs expand along both dimensions to tile the visual field (VF). **B)** Same as A for aspect ratio (AR) of 8. Note that at middle iterations, RFs form a one-dimensional rope that folds to tile the visual field. Final maps are qualitatively similar to those in Figure 3C. **C)** Greater aspect ratio (AR) produces striping of the azimuth map independent of the shape of the input visual field. Here a square input visual field results in the development of similar maps as those in Figure 3 with a circular visual hemi-field as input. **D)** Example single-cell RFs in V2, measured in anesthetized animal with two-photon calcium responses elicited by randomly flashed small squares covering the visual field (Methods). RFs are estimated using reverse correlations of stimulus frames with neural responses in the form of spikes extracted from calcium traces using Suite2p (*48*). ROIs are colored according to the azimuth value of the center of their RFs in the visual field. Some RFs, such as #1 and #2, exhibit slight elongations but their AR never reaches that of the cortical area (AR ∼ 8).

**Figure S4:**
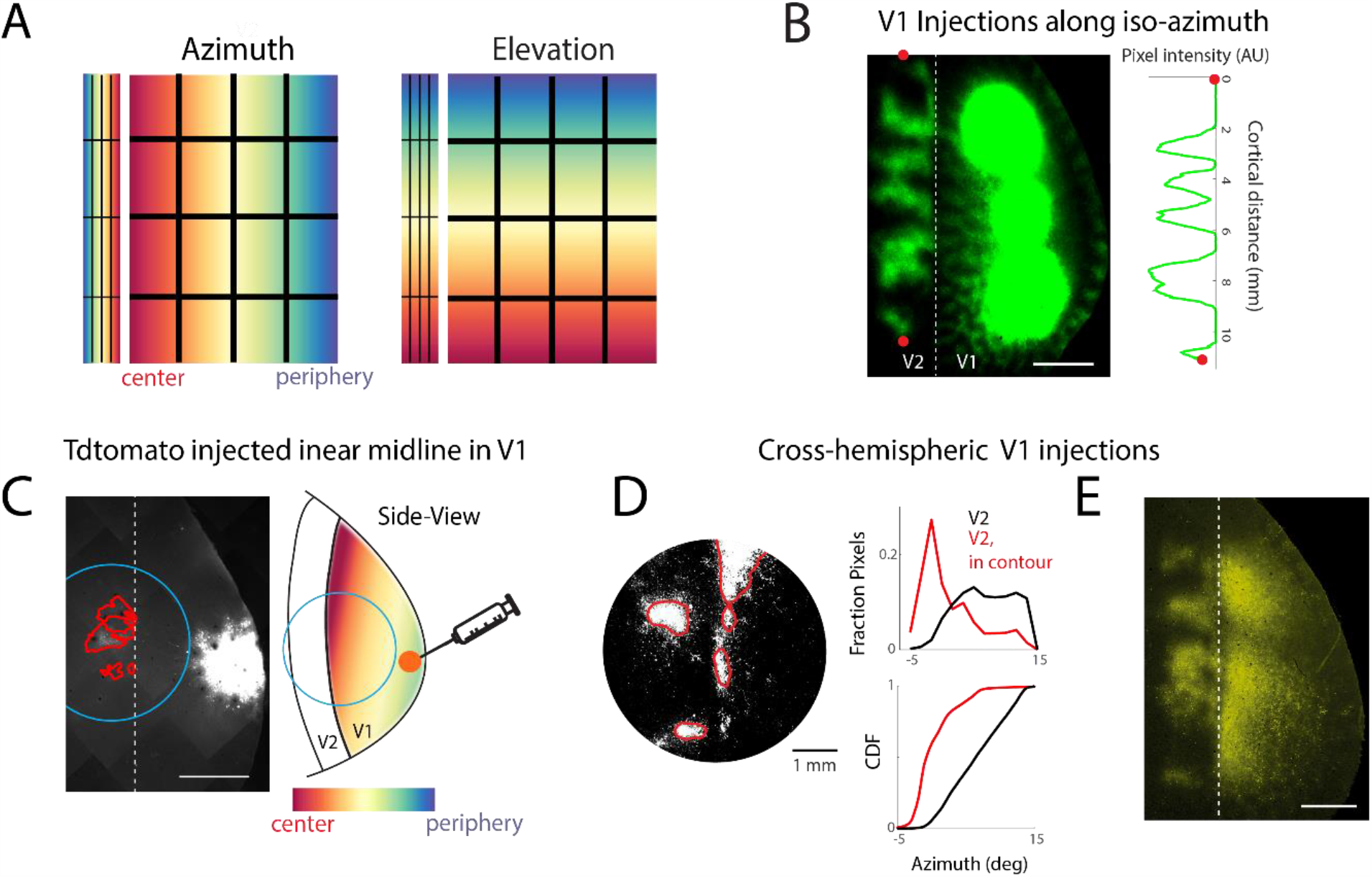
Supporting information for Figure 4. **A)** Relationship between maps in V1 and V2 under a linear transformation. Contrast with sinusoidal transform in Figure 4A. **B)** Injections of the same colored tracer along iso-azimuth axis in V1, produces a striped pattern of axons in V2, as evidenced by oscillations produced in the fluorescence intensity value along a linear track in V2. The gaps in V2 correspond to unlabeled central and peripheral regions in V1. **C)** Flattened section from left hemisphere of animal in Figure 4E showing site of V1 tracer injection outside of cranial window. **D)** Additional section from flattened left hemisphere for animal in Figure 4F showing callosal axon terminals at the border of V1 and V2 and in patches in V2 (left). Distribution of azimuth values for all pixels in V2, versus the pixels within the callosal axon terminal contours (right). The left-shift of the callosal distribution indicates the central visual field bias of these terminal patches. **E)** Callosal axon terminals produced in the right hemisphere from injections of tracer in left V1 near the V1/V2 border as shown in Figure 4C (middle). All flattened section scale bars are 2 mm.

**Figure S5.**
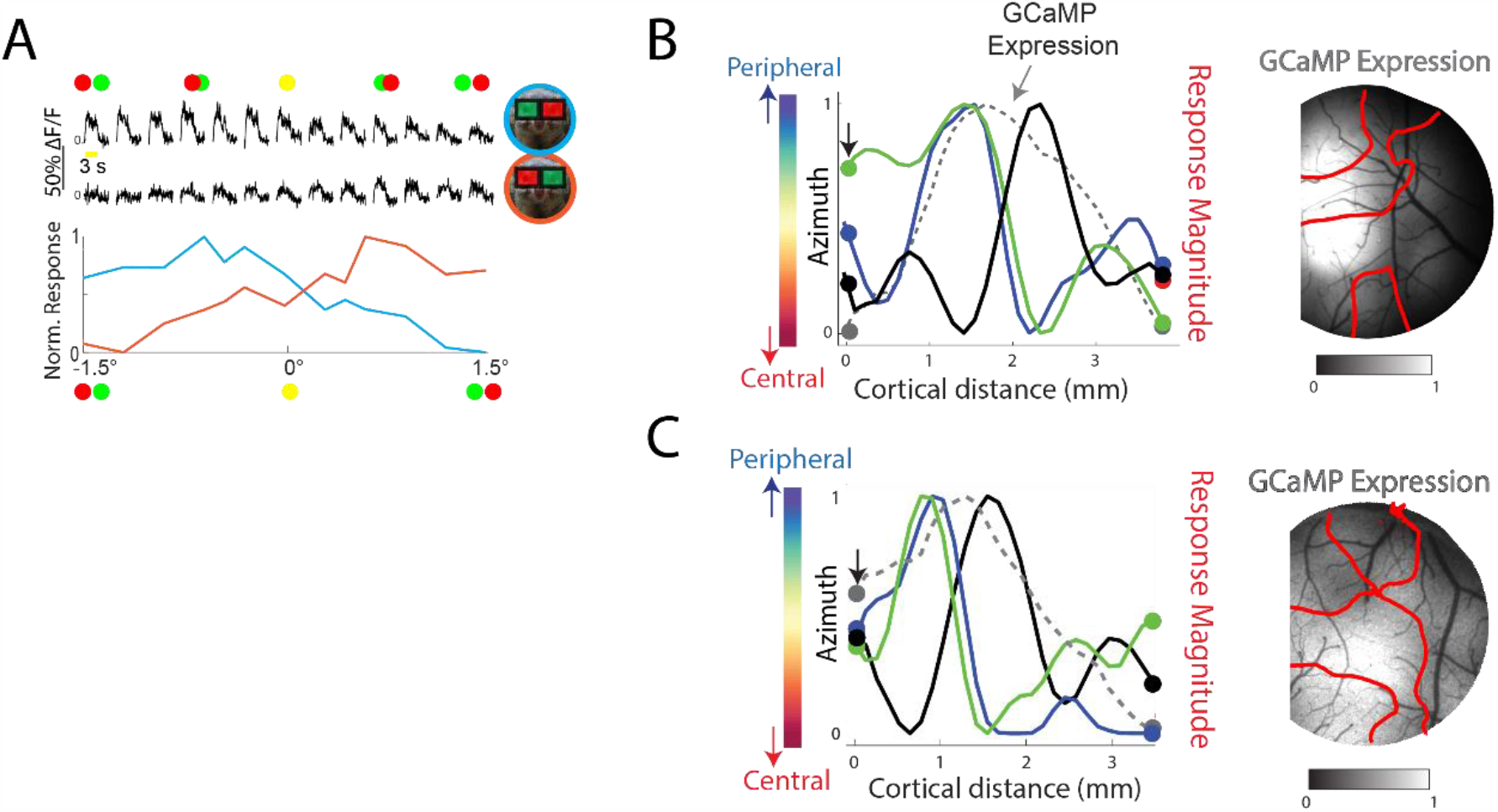
Supporting information for Figure 4. **A)** Two-photon fluorescence responses to random dot stereogram for disparity sensitivity. This cell prefers negative disparities (blue trace). When the red/green filter goggles are swapped the selectivity of the cell switches to positive disparities as expected. **B)** Same as Figure 5A, here overlaid with the extracted pixel values for GCaMP expression as a control (right). **C)** Same as B) for the animal in Figure 5B. GCaMP expression patterns are thresholded at mean +/-1SD.

## Supplementary Movies

Movies can be found on this YouTube playlist: https://www.youtube.com/playlist?list=PLhW0QP2TWaQiuyeu6d3yh0Q0U-NfNlRoP

**Movie S1. Retinotopy maps in tree shrew V1 and V2**. This video shows the elevation and azimuth maps in a 5 mm window placed over the border of V1 and V2. The maps were created by measuring GCaMP activity patterns in the window in response to bars moving along azimuth or elevation dimensions of the visual field. The pixels on the cortex are color-coded according to the position of the bar that elicited the largest response, on average over 30 trials. The right panel show the preferred visual field position for each corresponding region, indicated by the white moving box, on the cortex. The notable feature is that drawing a line on the cortex produces a linear path in the visual field in V1, but a sinusoidal path in V2. Thus the surface of V1 exhibits a linear transform of the visual field whereas the surface of V2 exhibits a sinusoidal transform.

**Movie S2. Sinusoidal transform of the visual field in V2**. This video shows the sinusoidal transformation between the visual field and V2. That is, it shows how regions in the visual field are mapped onto regions in V2. The colors corresponds to the azimuth (left/right) axis of the visual field. Then the colormap is changed to encode elevation, showing that the same transform also produces elevation maps similar to observed data. Since the map in V1 can be abstracted as a duplicate of the visual field for our purposes, this simulation can also be interpreted as showing the connectivity between V1 and V2. If one connects regions in V1 along the sinusoid, one gets the striped map in V2 that matches functional data.

**Movie S3. Sinusoidal transform of visual field in V2 - activation in V2 produced by bars in visual field**. This video simulates activity patterns in V2 in response to showing iso-azimuth or elevation bars in the visual field. This is similar to activating iso-azimuth/elevation regions of V1, which we simulate using tracer injections along iso-azimuth or iso-elevation regions in V1 (Figure 4). The gaps in activation patterns in V2 are due to inactivated regions along the sinusoidal path in V1/visual field.

